# Extreme sensitivity of fitness to environmental conditions; lessons from #1BigBatch

**DOI:** 10.1101/2022.08.25.505320

**Authors:** Kinsler, Schmidlin, Newell, Eder, Apodaca, Lam, Petrov, Geiler-Samerotte

## Abstract

The phrase “survival of the fittest” has become an iconic descriptor of how natural selection works. And yet, precisely measuring fitness, even for single-celled microbial populations growing in controlled laboratory conditions, remains a challenge. While numerous methods exist to perform these measurements, including recently developed methods utilizing DNA barcoding, all methods seem limited in their precision to differentiate strains with small fitness differences. This limit on precision is relevant in many fields, including the field of experimental evolution. In this study, we hone in on the factors that contribute to noisy fitness measurements and suggest solutions to avoid certain sources of noise. Surprisingly, even when common sources of technical noise are controlled for, we find that fitness measurements are still very noisy. Our data suggest that subtle environmental differences among replicates create substantial variation across fitness measurements. We conclude by providing best practices for obtaining precise fitness measurements and by discussing how these measurements should be interpreted given their extreme context dependence. This work was inspired by the scientific community who followed us and gave us tips as we live-tweeted a high-replicate fitness measurement experiment at #1BigBatch.

## Introduction

Measuring the relative fitness of mutant microbial strains has revolutionized our understanding of functional genomics and basic cell biology. For example, comparing the fitness of microbial strains with different gene deletions taught us about the function of thousands of genes (Giaever et al. 2002; Breslow et al. 2008), which genes work together (Costanzo et al.2016), and about the genetic programs that allow cells to respond to challenges like drugs, high temperature or nutrient deprivation (Gasch et al. 2000; Slavov and Botstein 2011; She and Jarosz 2018). Precisely quantifying the fitness of microbial populations is also of interest to evolutionary biologists, for example, those surveying the fitness effects of adaptive mutations (Levy et al. 2015; Venkataram et al. 2016; Kinsler et al. 2020), deleterious mutations (Wloch et al. 2001; Geiler-Samerotte et al. 2011; Johnson et al. 2019), or combinations of mutations (Flynn et al. 2020; Aggeli et al. 2021; Bakerlee et al. 2022). Further, measuring fitness of infectious microbial populations is of interest in evolutionary medicine (Brown et al. 2010; Nichol et al. 2019; Dunai et al. 2019), and measuring the fitness of engineered microbes is critical in industries focused on bioproduct production (Chubukov et al. 2016).

Given the importance of measuring microbial fitness in diverse fields, it is frustrating that we cannot measure fitness more precisely. Reported precision remains capped at detecting fitness differences on the order of 0.1 – 1% (Gallet et al. 2012; Leon et al. 2018; Duveau et al.2018)]. These estimates of precision may be inflated as they often ignore reproducibility issues that arise when performing replicate experiments on different days (Kinsler et al. 2020) or in different labs (Lithgow et al. 2017). On the other hand, natural selection can distinguish fitness differences that are orders of magnitude smaller than we can measure in the laboratory, depending on effective population size (Ohta 1973; Lynch and Conery 2003). This creates a problem for scientists: how do we understand the fitness effects of mutations if many of these effects are too small for us to measure?

### Methods to measure fitness and limitations on their precision

There are many reasons why the methods we use to measure fitness are limited in their precision. One common way to compare fitness across microbial strains is to measure each strain’s growth rate, in other words, the rate at which cells divide to make more cells. The change in cell density over time is often measured by the increase in optical density of liquid culture (Ram et al. 2019) or the increase in colony size on an agarose plate (She and Jarosz 2018) or a glass bottom plate (Levy et al. 2012; Sartori et al. 2021). This method has several limitations on precision, one being that measuring cell density is not as precise as counting individual cells, and another being that each microbial strain is usually grown separately and is separately affected by any biological or technical noise. A less common method to measure microbial growth rate is to track the level of a protein or transcript that controls growth, rather than measuring growth itself (Brauer et al. 2008; Geiler-Samerotte et al. 2013; Scott et al. 2014;Wu et al. 2022), though this method suffers the same limitations as those above.

Many consider the gold standard for measuring fitness to be an experiment where strains are competed in the same vessel (Hegreness et al. 2006; Kao and Sherlock 2008;Geiler-Samerotte et al. 2011; Ram et al. 2019). A benefit of this method over others is that all strains are grown simultaneously in the same well-mixed media in the same vessel and thus subject to the same exact environment. Further, fitness is defined more broadly than growth rate because it includes differential survival in conditions where growth is not proceeding exponentially or is halted (Ram et al. 2019). Finally, another benefit is that, in many implementations of this method, individual cells are counted, which is more precise than tracking changes in population density over time.

Some fitness competitions label competing strains with fluorescent proteins or genetic markers distinguishable in the presence of a drug or carbon source. Then, the fraction of cells labeled with each marker are counted as they change over time (Hegreness et al. 2006;Breslow et al. 2008; Kao and Sherlock 2008; Geiler-Samerotte et al. 2011; Gallet et al. 2012;Lenski 2017). These methods report percent error ranging from 0.01-1% and sources of noise including sampling noise, which worsens when fewer cells from each fraction are counted, and cell assignment noise, which worsens, for example, when fluorescent reporters are chosen that are more difficult to distinguish from each other (Gallet et al. 2012). More recent methods utilize “**DNA Barcodes**” to differentiate strains (Levy et al. 2015; Venkataram et al. 2016; Kinsler et al. 2020; Bakerlee et al. 2021). These are short regions of the genome that are unique to a given strain and flanked by consensus sequences such that all barcodes can be amplified using the same primer pair. Next generation sequencing allows for tracking the frequency of each DNA barcode over time with incredibly high throughput, such that millions of individual cell barcodes can be counted at each timepoint. Fitness estimates are obtained by quantifying how the barcode that labels a particular strain changes in frequency over time relative to a reference (**Fig 1A**). In barcoded fitness competitions, as in competitions with fluorescent markers, sources of noise include sampling noise, which can be reduced by sequencing barcodes at high coverage at every timepoint. Introducing a larger number of replicate experiments is another common way to decrease the reported error on fitness, though this assumes fitness is insensitive to subtle environmental differences between replicates.

**Figure 1:**
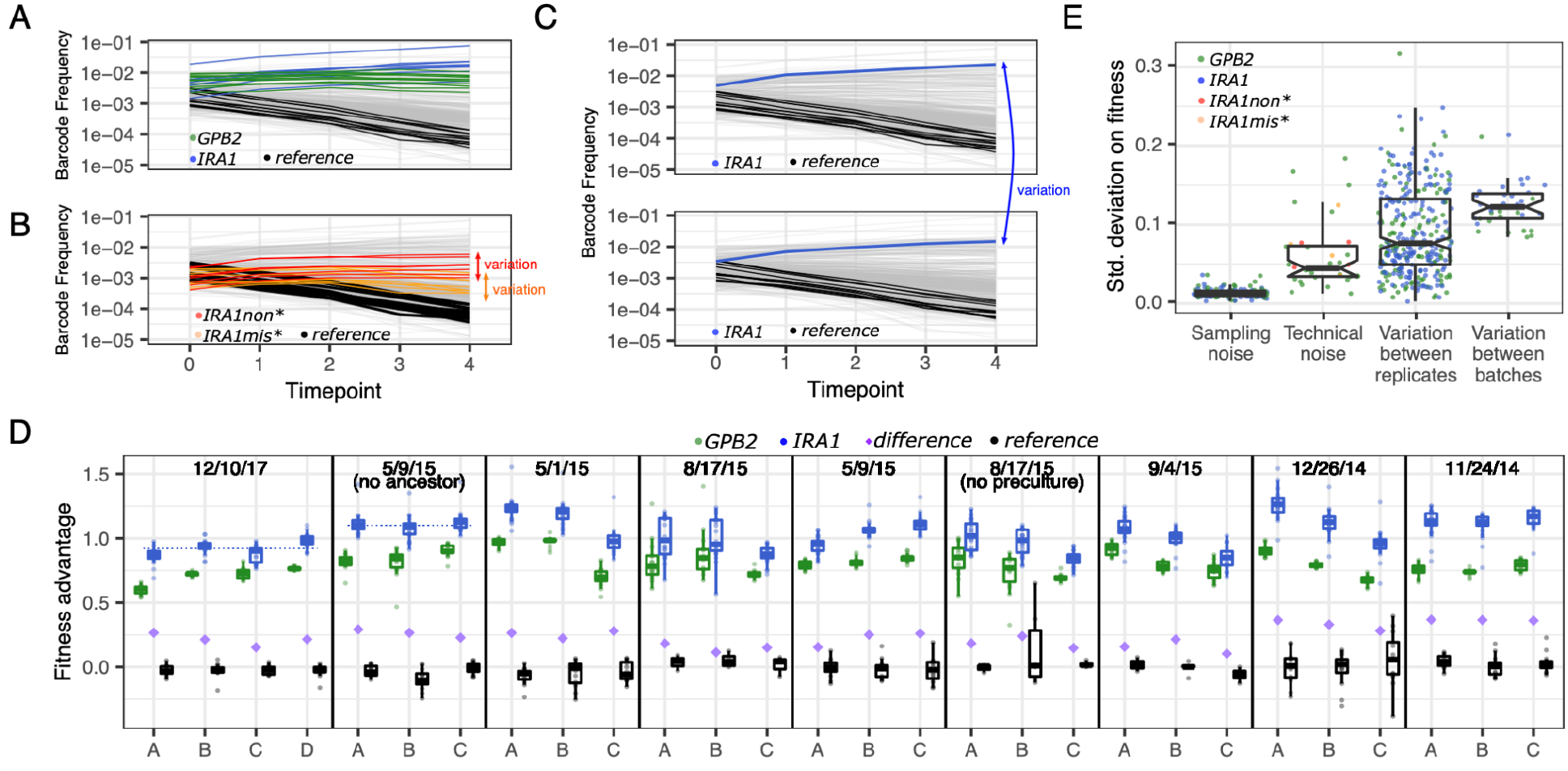
How fitness is estimated and how these estimates vary. **(A)** These example barcode frequency trajectories show the raw data used to estimate fitness for hundreds of barcoded lineages competing within a single vessel. Trajectories for about 500 barcoded lineages are shown (grey) with 9 lineages each possessing a unique mutation to the *IRA1* gene (blue), 13 lineages each possessing a unique mutation to *GPB2* (green), and 13 uniquely barcoded reference lineages lacking mutations (black). Fitness is inferred from the change in barcode frequency over time relative to the average rate of change of the reference lineages. **(B)** This panel clarifies how we study technical variation by observing the fitness of identical genotypes competing within the same vessel. We studied 6 strains that each possess a frameshift AT to ATT insertion at bp 4090 in *IRA1* (red’) as well as 8 strains that each possess a G to T mutation at bp 3776 in *IRA1* (orange). **(C)** This panel clarifies how we study variation in fitness across batches and replicate experiments. To do so, we calculate how much variation exists in fitness estimates for a single unique barcode when measured in different experiments. **(D)** Each panel represents 3 to 4 replicate fitness competition experiments (labeled A, B, C, or D) that were performed in the same “batch” (i.e., on the same date following the same protocol). Minor batch-specific protocol modifications are listed in parentheses. Each point represents the fitness of a uniquely barcoded yeast lineage with a mutation in the *IRA1* (blue) or *GPB2* (green) gene, or a reference strain possessing no mutations (black). Boxplots summarize the distribution across all lineages possessing mutations to the same gene, displaying the median (center line), interquartile range (IQR) (upper and lower hinges), and highest value within 1.5 × IQR (whiskers). Purple diamonds represent the average fitness difference, per replicate, between all lineages possessing *IRA1* vs. *GPB2* mutations. Dotted blue lines guide the eye to observe a “batch effect,” whereby the fitness of lineages possessing nonsense mutations in the *IRA1* gene differs across experiments performed on different days. **(E)** Each point represents the standard deviation across multiple fitness measurements. For each category listed on the horizontal axis, boxplots summarize the same features as **panel D**, with notches representing a roughly 95% confidence interval around the median calculated as 1.58 × IQR / √n. The points pertaining to each category on the horizontal axis were calculated as follows: *Sampling noise:* We simulate sampling noise by randomly resampling barcodes with replacement from all time points of a single experiment, to the same coverage, and re-calculating fitness. The standard deviation across 5 samples is reported for 9 *IRA1* lineages and 13 *GPB2* lineages (22 observations). *Technical noise:* We calculate the standard deviation on fitness within a single experiment for all lineages carrying any mutation in *GPB2* (green), carrying an AT to ATT insertion at bp 4090 of IRA1 (red), or carrying a G to T mutation at bp 3776 of *IRA1* (orange). We do this for each of 28 replicate experiments in which multiple *GPB2* lineages are present, and 4 experiments in which multiple IRA1non* or IRA1mis* lineages are present (36 observations). *Between replicates:* We calculate the standard deviation on fitness across multiple experiments performed on the same day (i.e., in the same batch). The fitness of 9 barcoded lineages possessing mutations to *IRA1* was calculated in 9 separate batches, as was the fitness of 13 uniquely barcoded lineages with mutations to *GPB2* (9×9 + 13×9 = 198 observations). *Between batches:*We calculate the standard deviation on fitness across all 28 replicate experiments. The fitness of 9 barcoded lineages possessing mutations to *IRA1* was calculated, as was the fitness of 13 uniquely barcoded lineages with mutations to *GPB2* (22 observations).

### Even with high sequence coverage and replication, we observe noisy fitness data

In our study, we quantify noise in barcoded fitness competitions in *S. cerevisiae*, focusing on barcodes that received extremely high sequencing coverage to minimize sampling noise and that were studied across 28 replicate experiments. We found surprisingly high noise, for example, certain adaptive mutants have fitness advantages that range from 180% of the reference in one replicate experiment to 220% in others. We performed follow-up experiments to identify the source of this variation, including 79 technical replicates in which we re-analyze the data from a single competition experiment using different protocols. Previous work has suggested that stochastic events during barcode extraction and amplification may be the largest contributor to noise for barcodes that received high sequencing coverage (Levy et al. 2015; Venkataram et al. 2016). However, we did not find that these sources contribute strongly to variation in fitness when we followed the extraction and amplification protocols recommended by previous studies. While we did find one major source of technical noise, index mis-assignment during sequencing, the dominant source of noise in our experiments is not necessarily technical. Instead we observe large “batch effects”, i.e. differences that arise between ostensibly identical experiments performed on different days. These batch effects may reflect biologically relevant but subtle changes in environmental conditions that influence fitness, for example, slight differences in temperature, media composition, humidity, shape of the culture vessel, shaking speed, etc.

This observation that fitness may be extremely sensitive to environmental context makes us question how to obtain and interpret fitness measurements. Are studies, such as ours, that investigate the best practices for reducing technical sources of noise still worthwhile? Should we instead focus on minimizing replicate and batch effects, perhaps by using incubators with extremely low variation in temperature or ultra precise scales for media preparation? Or should we embrace variation in fitness and search for new insights about genotype-by-environment interactions by studying the way fitness varies across replicates and batches (Kinsler et al.2020)? We argue that all these approaches have value and report a complete list of ideas to improve the way we obtain and interpret fitness measurements at the end of our manuscript. We hope the findings of our endeavor to generate ultra precise fitness measurements, which we live-tweeted about at #1BigBatch (KerryGeilerSamerotte 2017; Kinsler 2017), will act as a user’s guide for others quantifying fitness by utilizing DNA barcodes, will inspire important questions about how to measure and report fitness given its sensitivity to environmental changes, and will fuel the larger discussion on how to conduct reproducible and precise laboratory studies.

## Results

### Experimental design to distinguish sources of noise in barcoded fitness competitions

Our goal is to dissect the sources of noise that contribute to variation in fitness beyond sources that are easy to understand such as the sampling noise that is introduced when barcodes are sequenced at low coverage. We start by focusing on variation in fitness for barcoded mutants that had been sequenced at extremely high coverage. We also focus on strongly adaptive mutants as these increase in frequency over the course of a competition experiment, thus sequencing coverage continues to improve over time (**Fig 1A**). Previous experimental evolutions utilizing DNA barcodes discovered two types of strongly adaptive mutants that commonly arise in baker’s yeast in response to glucose limitation: nonsense mutations to the *IRA1* gene and missense mutations to *GPB2* (Levy et al. 2015; Venkataram et al. 2016). The fitness of these mutants in glucose-limited conditions has been measured 28 times using barcoded fitness competitions (Kinsler et al. 2020). In these 28 replicate experiments, the average coverage of a barcoded lineage possessing an *IRA1* nonsense mutation was 37,690 reads per timepoint. For lineages possessing mutation in *GPB2*, average coverage per timepoint was also very high (13,321 reads). For comparison, previous barcoded fitness competitions aim for coverage thresholds that are smaller by orders of magnitude (100 to 200 reads per mutant per timepoint) (Venkataram et al. 2016; Kinsler et al. 2020).

Focusing on fitness variation that is unlikely to be due to sequencing sampling noise allows us to distinguish between other sources of noise. First, we wanted to disentangle biological noise that reflects environmental differences across replicate experiments from technical noise that reflects stochastic events that take place when barcodes are prepared for sequencing. Technical noise includes variation in fitness introduced when DNA is extracted from cells, when barcodes are amplified via the polymerase chain reaction, and when samples are multiplexed prior to sequencing. Later in this study we describe experiments that isolate each of these sources of technical noise, but first we compared the overall amount of noise arising from technical vs. biological sources (**Fig 1**). To do so, we compared variation in fitness of identical mutants competing within the same vessel (**Fig 1B**) to variation in fitness of identical mutants across replicate competition experiments (**Fig 1C**). Since mutants competing within the same vessel experience the same environment, we assumed that variation in their fitness reflects technical and not biological noise (**Fig 1B**). To study identical mutants competing within the same vessel, we utilized strains that were engineered to possess identical mutations in the *IRA1* gene, but were given different barcodes (Kinsler et al. 2020). We studied 6 such strains that each possess a frameshift AT to ATT insertion at bp 4090 in *IRA1* (referred to herein as ‘*IRA1*non*’) as well as 8 such strains that each possess a G to T mutation at bp 3776 in *IRA1* (called *‘IRA1*mis*’) (**Fig 1B**). We also utilized strains with adaptive mutations in *GPB2*, because previous work found that different mutations to this gene had similar effects on fitness, regardless of their position within the open reading frame (Levy et al. 2015; Venkataram et al. 2016). Quantifying variation in the fitness of identical (*IRA1*mis* and *IRA1*non*) or similar (*GPB2*) strains when they are competing within the same vessel sets an upper limit on the amount of noise that comes from technical sources. We call this an upper limit because some of the noise might still reflect biological variation, for example, because of microenvironmental differences within a culture vessel.

In addition to quantifying the upper limit on technical noise, we also sought to quantify biological noise by studying how fitness measurements vary across 28 replicate fitness competition experiments. To do so, we quantified how the fitness of a unique barcode, marking a unique mutation, varies between experiments (**Fig 1C**). These comparisons (**Fig 1C**) tell us about the extent to which additional sources of noise, on top of those observed within a single competition experiment (**Fig 1B**), contribute to variation in fitness.

### We observe more variation in fitness than can be explained by sampling noise

Across all 28 replicate experiments, we observe that nonsense mutations to *IRA1* and mutations to *GPB2* are always adaptive, meaning they always have higher fitness than a reference strain lacking any of these mutations (**Fig 1D**; blue and green boxplots are always above black boxplots). However, there is substantial variation in their fitness advantage relative to the reference strain. For example, the median fitness across all barcoded lineages possessing a nonsense mutation in the *IRA1* gene varies across replicates from 180% of the reference fitness to 220% (**Fig 1D**; the height of the blue boxplot varies). What causes this variation in fitness?

Previous work suggests that sampling noise is an important source of variation in barcoded fitness competitions (Levy et al. 2015; Venkataram et al. 2016). However, as mentioned above, we chose to focus on *IRA1* and *GPB2* mutants that received high sequencing coverage and for which sampling noise should be low. Further, we estimated the impact of sampling noise on fitness estimates by randomly resampling from the barcode frequency data while preserving the same level of coverage. We find that sampling noise is insufficient to explain the amount of variation in fitness that we see (**Fig 1E**; leftmost boxplot).

### Technical noise influences fitness measurements

Our fitness inference method yields different fitness estimates for identical mutants competing against one-another in the same vessel (**Fig 1B**). Barcoded mutants possessing identical mutations (*IRA1*mis* and *IRA1*non*) were included in each of the four replicate experiments performed on 12/10/17. Variation within each experiment in the fitness of the *IRA1*mis* and *IRA1*non* lineages is too large to be explained by sampling noise (**Fig 1E**; red and orange points in second boxplot from left). In addition, all 28 replicate experiments each include 13 lineages possessing different mutations to the *GPB2* gene. On average, variation in the fitness of *GPB2* mutants competing within the same vessel tended to be greater than sampling noise (**Fig 1E**; green points in second boxplot from left). Thus, we conclude that technical noise beyond sampling noise must affect fitness estimates. We explore these technical sources of noise later in this manuscript.

### We observe additional variation in fitness across replicates and batches

Next we sought to address the question of whether additional sources of biological noise, on top of the technical noise observed between identical mutants competing within a single experiment (**Fig 1B**), contribute to variation in fitness across replicate experiments (**Fig 1C**). Across all replicate experiments, and more so across replicates performed on different days (i.e., in different batches), we see additional variation in fitness beyond what we observed among identical mutants competing within the same vessel (**Fig 1E**; rightmost two boxplots). These differences are substantial enough that they are visible in the fitness data (**Fig 1D**). For example, in the batch of fitness competitions performed on 12/10/17, the average fitness advantage of *IRA1* mutations is significantly smaller than in the batch of competitions performed on 5/9/15 (**Fig 1D**; compare panels 1 v. 2). And in the 3 replicate fitness competitions performed on 5/1/15, the average advantage of *IRA1* nonsense mutations differs, with this advantage being significantly lower in replicate ‘C’ (**Fig 1D**; compare replicates within panel 3).

Given the largest contributor to noise appears to be batch effects (**Fig 1E**), we wondered, what are the batch effects and how can we avoid them to get more precise measurements of fitness? Several studies suggest that differences across batches, or even across replicates, are not caused by the same sources of technical noise that influence individual experiments (Lithgow et al. 2017; Kinsler et al. 2020). Instead, these differences may reflect what we term ‘biological noise’ arising when fitness is sensitive to subtle differences across environmental conditions. Such environmental differences may include variation in the preparation or age of the media, fluctuations in the temperature or humidity of the incubator, or stochastic differences in the composition of the starting pool of barcoded lineages (Kinsler et al.2020). Previous work was able to predict fitness variation across batches by modeling how mutant fitness changes across subtle environmental perturbations. Further, the variation across batches was informative about fitness variation across very different environments (Kinsler et al.2020). This provides support for the idea that batch effects represent subtle genotype-by-environment interactions, rather than random technical noise. The strong contribution that batch effects make on variation in fitness highlights the importance of a careful experimental design where the effects of treatment are not confounded with the effects of batch (**Box 1**). It also highlights how strongly the impact of mutations depends on context, including even subtle contextual changes that are present across batches (Eguchi et al. 2019;Geiler-Samerotte et al. 2020; Kinsler et al. 2020).

#### Box 1 - Improving precision by adding more replicates can fail due to batch effects

Improving fitness measurement precision by adding more replicates makes several assumptions. For example, this method assumes that between-replicate (or between-batch) variation is uncorrelated technical noise, rather than variation in fitness due to environmental differences. For example, consider a case where fitness has been measured in a control condition and in the presence of a drug (**Fig B1**). If we fail to consider the effect of hidden variables that vary from batch to batch, then in this case we would correctly infer that the fitness is higher in the control condition when we take the average across all replicates and batches (**Fig B1**; rightmost ‘aggregate’ panel). Our confidence in the precision of our fitness measurements (as measured by standard error - 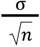), would be fairly high, given the relatively large number of replicates across all batches. However, if we had measured fitness in the drug condition only in batch one, and the control condition only in batch two (**Fig B1**; dotted box), we would have incorrectly observed that the drug has no effect on fitness. In this case as well, our confidence in this fitness measurement would be fairly high, given we performed 3 replicates for each condition. In both cases, our confidence in the measurement is overstated, given the standard error may be over-representing how precisely fitness can be measured, as there are uncontrolled factors that vary from batch to batch that change the fitness effects of the mutant across the environments (**Fig B1**).

**Figure B1.**
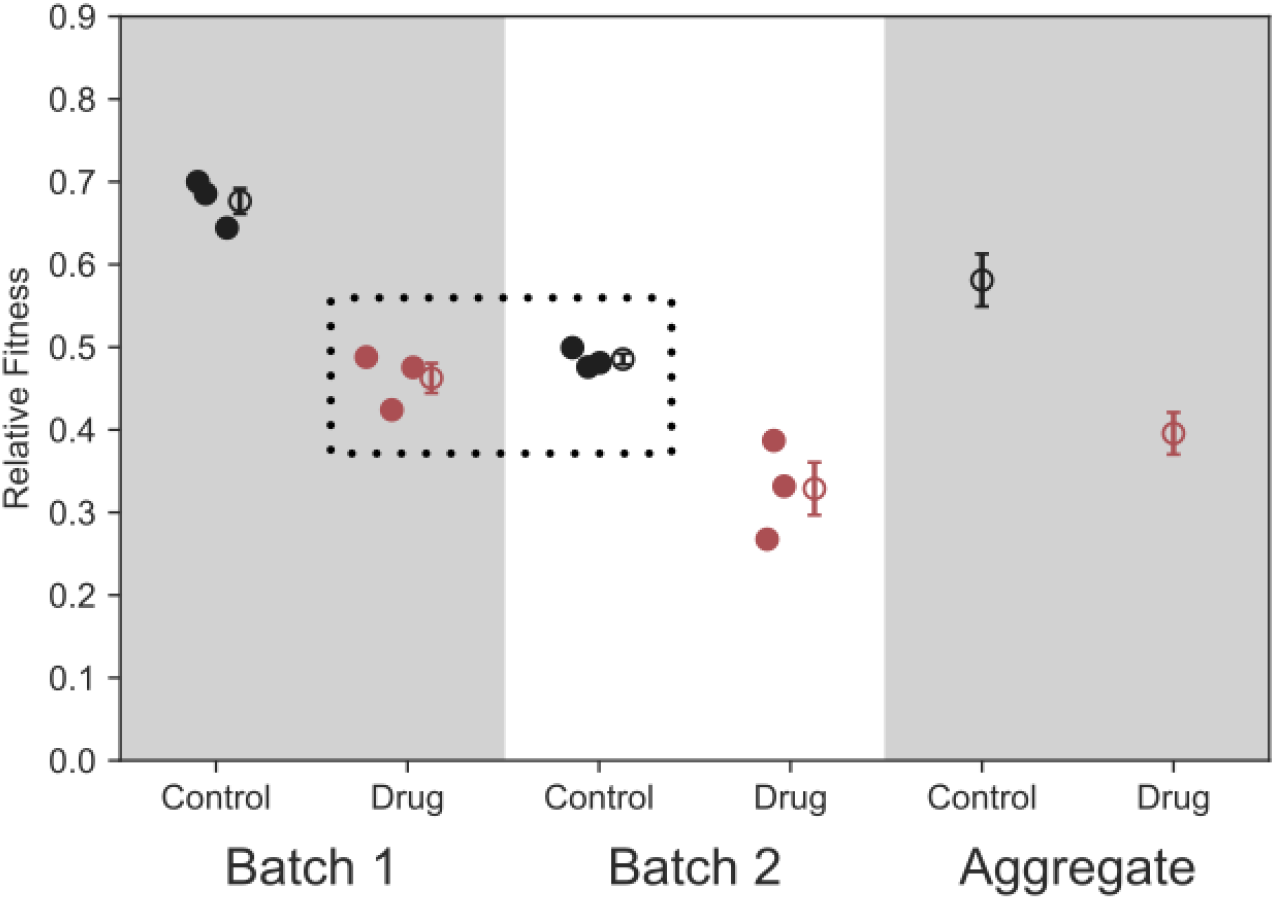
Problems introduced by batch effects. This figure shows a toy example using simulated data that reflects how aggregating across measurements from different batches (e.g., measurements performed on different days) can ignore batch-to-batch variation leading to an overestimate of measurement precision. Each of the 3 dots represents replicate fitness measurements of a single genotype. In this example, batch effects are at least as strong as the effect of the drug on fitness. Since both drug and control conditions were included in each batch, the aggregate panel accurately captures the fitness difference between the drug and control, but overestimates the precision of this measurement. Had the experiment been designed differently, with drug and control conditions being surveyed on different days (dotted box), the effect of the drug on fitness may have been obscured by batch effects.

In addition to environmental differences between batches, another potential source of batch effects may be the inference method used to calculate the fitness of each mutant from changes in barcode frequency over time (Venkataram et al. 2016; Li et al. 2018a; Kinsler et al.2020). Though there appears to be large variation in the fitness of IRA1 nonsense mutants across replicates and batches (**Fig 1D**; blue), the rank order of fitness appears to be conserved such that *IRA1* nonsense mutants are always more fit than *GPB2* mutants (**Fig 1D**; blue boxplots are always higher than green). Indeed, the fitness difference between *IRA1* and *GPB2* mutants (**Fig 1D**; purple diamonds) shows reduced variation across replicates relative to the individual fitness estimates for each type of mutant (**Fig 1D**; std dev across purple diamonds is 0.032, while std. dev. across boxplot medians is 0.117 for *IRA1* and 0.09 for *GPB2*). This may indicate that differences between replicates and batches are in part not due to differences between experimental conditions and are instead introduced during the fitness inference procedure. One key step of the fitness inference method determines either the mean fitness of the population (Venkataram et al. 2016; Li et al. 2018a) or the fitness of spiked-in reference strains (Kinsler et al. 2020). Then, it sets all other fitnesses by these initial baseline inferences (**Box 2**). If these initial inferences differ across replicates or batches, it could lead to variation in inferred fitness values that preserves the rank order of mutant fitness, as observed in **figure 1D**. Thus, another way to improve precision in fitness estimation may be to improve the accuracy of these initial inferences, for example, by spiking in a larger number of reference strains, which serve as a baseline for estimating the impact of adaptive mutations on fitness.

#### Box 2 - Inference of a fitness benchmark is critical to reproducible measurements

During a fitness competition experiment, a strain’s change in frequency depends on its fitness relative to the fitness of the other strains in the population (Li et al. 2018a). Thus, benchmarking the fitness of a population is a crucial step to properly normalizing fitness measurements and ensuring reproducibility across environments and batches. There are two common methods to calculate a fitness benchmark. One method directly calculates the mean fitness of an entire population by using the inferred fitnesses and frequencies of strains in the population (Li et al. 2018a). This method is commonly used in screens where many mutations are being simultaneously assayed (Fowler and Fields 2014;Sharon et al. 2018; Liu et al. 2020), as it’s assumed that most mutations have little to no effect on fitness such that this average is representative of the fitness of an unmutated strain. Another method for benchmarking fitness is the use of reference strains. With this approach, a reference strain/s is spiked into the population, and fitnesses of other strains are explicitly calculated relative to this reference. This approach allows fitnesses to be easily compared across environments, because they are always relative to the same reference strain. When studying the fitness of adaptive mutations, the reference strain is often the unmutated ancestor of an evolution experiment (Venkataram et al. 2016; Kinsler et al. 2020). Given this ancestor will often fall in frequency as the adaptive mutants rise in frequency, it is advisable to spike it in a high concentration to ensure adequate coverage (and low sampling noise) and to include many uniquely barcoded copies to contend with variation in fitness arising from technical sources.

However, differences in mean fitness inference are unlikely to entirely explain batch effects. If they did, we would expect to see the same amount of variation across replicates and batches. However, variation across batches is significantly greater (**Fig 1E**), suggesting environmental differences between batches may contribute. Also, the observation that the rank order of fitness is conserved across batches is not necessarily indicative of differences in mean fitness inference. Subtle environmental differences between batches may have a similar effect on the fitness advantage of both *IRA1* and *GPB2* mutants, thus preserving their rank order. Finally, previous work demonstrates that the batch effects we study here indeed reflect subtle genotype-by-environment interactions (Kinsler et al. 2020).

In sum, batch effects, and to a lesser degree, replicate effects, appear to be prevalent in barcoded fitness measurements (**Fig 1D & E**). They are likely caused by a combination of biological sources (e.g., subtle differences in the media, equipment, strain composition, and/or protocol) and technical sources (e.g., variation in the fitness inference procedure). On one hand, batch effects may represent a latent opportunity to learn new biology, e.g., which mutants are most sensitive to which environmental perturbations. On the other hand, batch effects can be extremely problematic in some circumstances. For example, they can make typically reported measures of precision, such as percent error across replicates, difficult to interpret (**Box 1**). And they can unfairly exaggerate fitness differences between environments that were studied on different days, or conversely, they can obscure these differences (**Box 1**). The severity of the problem depends on the magnitude of the fitness differences that the researcher is hoping to capture. In the field of experimental evolution, where researchers are often interested in understanding subtle fitness differences across environments (Venkataram et al. 2016; Li et al.2018b, 2019; Jerison et al. 2020; Kinsler et al. 2020; Boyer et al. 2021; Bakerlee et al. 2021), batch effects may be a large problem.

If each replicate, and to a greater extent, each batch experiment, measures fitness in a different environment, then we cannot improve fitness precision in a single environment by adding more replicates. How, then, can we improve the precision of fitness estimates? To answer this question, we performed additional experiments to explore three potential sources of technical noise that may limit precision on fitness estimation within a single experiment: 1) inefficient DNA extraction, 2) stochastic events during barcode amplification, and 3) index mis-assignment during barcode sequencing.

### Stochastic events during barcode extraction and amplification contribute little noise

In order to infer fitness from changes in barcode frequency over time during a fitness competition experiment (**Fig 1A**), one must first extract DNA from a large number of cells at many timepoints (KerryGeilerSamerotte 2018a). Next, one must amplify the barcode region and prepare it for sequencing by attaching multiplexing indices and illumina indices. Both of these steps can introduce imprecision into a fitness measurement. For example, if DNA is extracted from too few cells, or extracted in a biased way such that cells with certain mutations contribute an unfair quantity of DNA, fitness estimates may be imprecise or skewed. Similarly, if PCR is inefficient such that only a small number of barcodes are amplified in the early cycles, these barcodes will become overrepresented in a way that does not reflect their true abundance (i.e., PCR jackpotting). To understand the contribution of these two sources of imprecision, we performed dozens of technical replicate experiments (KerryGeilerSamerotte 2018b; Kinsler 2018a) where we split cells from a single timepoint and either performed multiple replicate PCRs (**Fig 2A**) or multiple replicate DNA extractions (**Fig 2B**). Then, we compared the barcode frequencies from pairs of technical replicates to see how much noise (measured via R^2^) is introduced by PCR and by DNA extraction. Previous work suggested that these two sources of noise contribute to imprecise fitness measurements for highly adaptive mutants like the ones we study here (Levy et al. 2015; Venkataram et al. 2016; KerryGeilerSamerotte 2018c).

**Figure 2:**
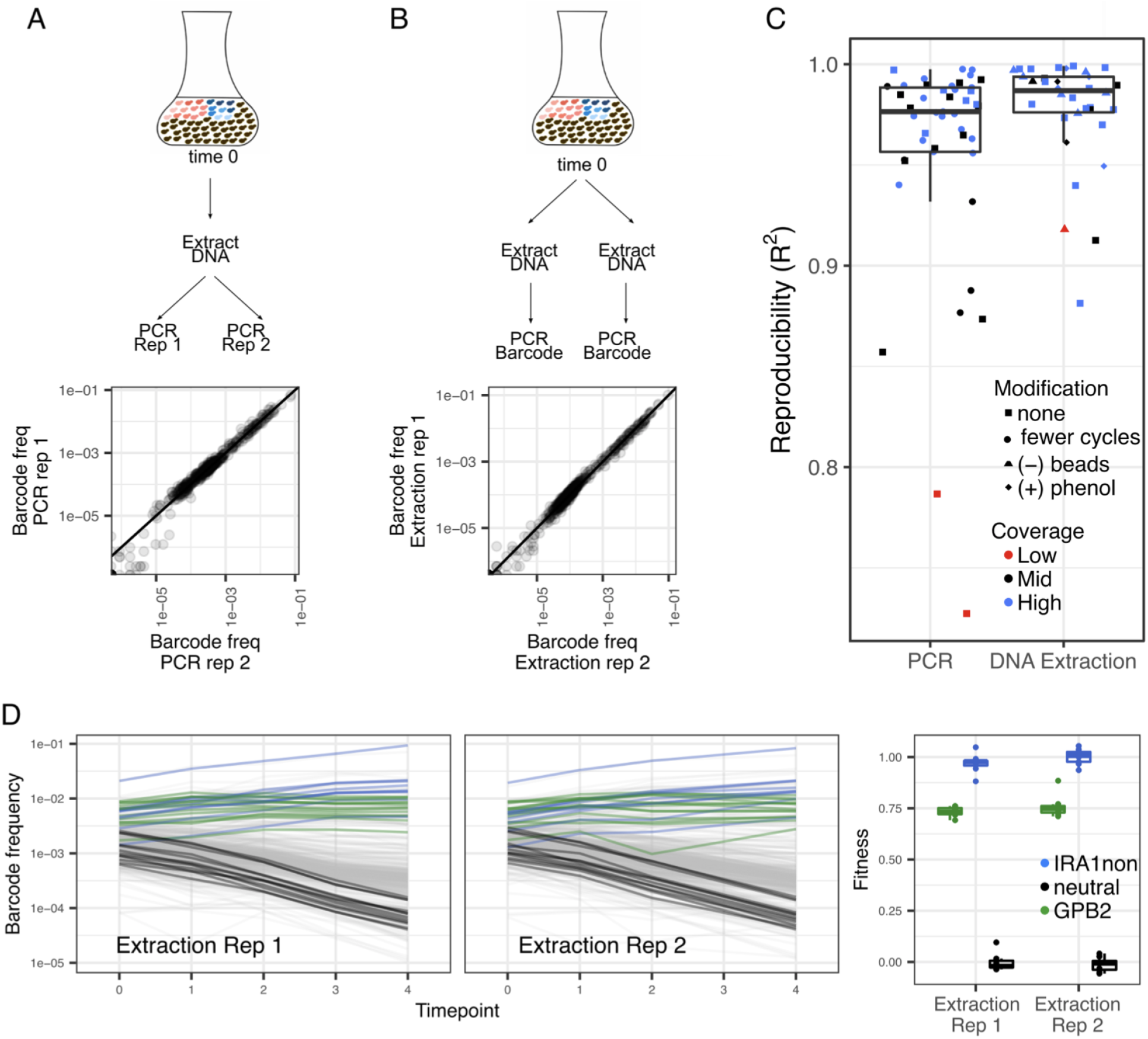
Stochastic noise generated during barcode extraction and amplification appear to contribute little noise to fitness estimation. **(A)** Schematic showing how we divided a sample to perform a PCR replicate and comparison of barcode frequencies for a pair of replicates where every point represents one of 500 barcodes. **(B)** Schematic showing how we divided a sample to perform an extraction replicate and comparison of barcode frequencies for a pair of replicates where every point represents one of 500 barcodes. **(C)** Reproducibility (R^2^) is greater than 0.9 for most pairs of PCR and extraction replicates. Modifications to the PCR or extraction procedure did not seem to affect reproducibility, though low sequencing coverage did often result in lower reproducibility. Boxplots summarize the distribution across all replicates of the same type, displaying the median (center line), interquartile range (IQR) (upper and lower hinges), and highest value within 1.5 × IQR (whiskers). **(D)** Frequency trajectories look similar across two example technical replicates, as do inferred fitnesses of representative mutants. Boxplots summarize the distribution across all mutant lineages of the same type, displaying the median (center line), interquartile range (IQR) (upper and lower hinges), and highest value within 1.5 × IQR (whiskers).

We found that stochasticity during PCR amplification contributes only a small amount of noise in our experiment (**Fig 2C**). In total, we compared 45 pairs of technical replicate PCRs. In some of these, both replicates included 27 PCR cycles; in others one replicate included fewer total cycles (22 v. 27). These deviations in cycle number did not appear to lead to greater imprecision. The typically high reproducibility we saw between PCR technical replicates could indicate that our PCR protocol is robust to effects like PCR jackpotting. In brief, we utilized a two-step PCR approach. The first step tags molecules with a unique molecular index (UMI) to differentiate uniquely sampled barcodes from barcodes that were lucky and were duplicated multiple times during the second step of PCR. We also split each PCR into several tubes such that if PCR jackpotting occurs, it will not occur for the same barcodes in every tube (Levy et al.2015; Venkataram et al. 2016; Kinsler et al. 2020). For a few of our technical replicates, we observed lower reproducibility (R^2^ < 0.9 for 6/45 pairs of PCR replicates). This lower reproducibility seems more common in cases where one replicate of the pair received low (on average <20 sequencing reads per barcode) or moderate (on average 20 – 600 sequencing reads per barcode) coverage (**Fig 2C**).

We found that stochasticity during DNA extraction also contributes only a small amount of noise (**Fig 2C**). In total, we compared 34 pairs of technical replicate DNA extractions, some of which were performed following protocols that involved harsher treatments (e.g., bombarding cells with glass beads or adding phenol to help them release their DNA). These modifications did not appear to affect reproducibility (**Fig 2C**). The high reproducibility we saw between technical replicates could indicate that our protocol is effective at extracting DNA from a very large number of cells in an unbiased way (Kinsler et al. 2020). An important note is that any noise detected from DNA extraction replicates also includes noise contributed during the PCR step (**Fig 2B**). Thus, we have de facto performed 79 (45+34) technical replicate PCRs, with 72/79 (91%) having reproducibility greater than R^2^ = 0.9.

Since DNA extraction and PCR amplification are performed at every timepoint, we wanted to understand how the aggregate noise introduced across many timepoints by these procedures would affect fitness estimates. To do so we studied frequency trajectories from the experiment where we performed DNA and PCR technical replicates on the largest number of timepoints (4 out of 5 timepoints) (**Fig 2D**). We found very similar fitness measurements were inferred from the barcode frequency trajectories comprised of each set of technical replicates. This suggests that, when following the protocols we use here, developed in earlier work (Levy et al. 2015; Venkataram et al. 2016; Kinsler et al. 2020), noise from PCR amplification and DNA extraction has a minor effect on fitness precision (**Fig 2D**). Thus, we continued to explore other sources of noise that might affect barcoded fitness competitions.

### Template switching on patterned flow cells appears to be a major source of noise

We found that template switching is a major source of noise in barcoded fitness competitions, and also discovered best practices for dramatically reducing this problem. The 28 experiments in **figure 1** were performed using these best practices, thus template switching cannot explain the variation in fitness that we observe in **figure 1**. Below we define template switchingg, explain our best practices to contend with it, and describe a series of experiments designed to understand its extent and mechanism.

In order to combine several samples on a single lane of sequencing, researchers often make use of index primers that label each sample on the lane. In the past, combinatorial indexing schemes have been used to identify samples such that each pair of indices uniquely labels a sample (**Fig 3A**). However, previous work, with relatively diverse RNASeq libraries, has shown platforms that utilize Illumina’s ExAmp chemistry (e.g., HiSeq 4000, HiSeqX, NovaSeq) have substantial rates of mis-assigned indices, such that 5-10% of sequencing reads are mis-assigned (Sinha et al.). This means that a substantial number of reads from one sample can appear to belong to another sample, causing errors with downstream analysis. These errors are especially problematic for analyses relying on quantitative measurements of abundances (e.g., barcode sequencing, RNASeq, or other amplicon-based sequencing such as 16S, deep mutational scanning, and massively-parallel reporter assays). Unfortunately, most previous studies of index mis-assignment have focused on RNASeq, rather than barcode sequencing. Here we designed an experiment to understand the frequency of index mis-assignment in barcode sequencing, as well as the underlying mechanisms and potential solutions.

**Figure 3.**
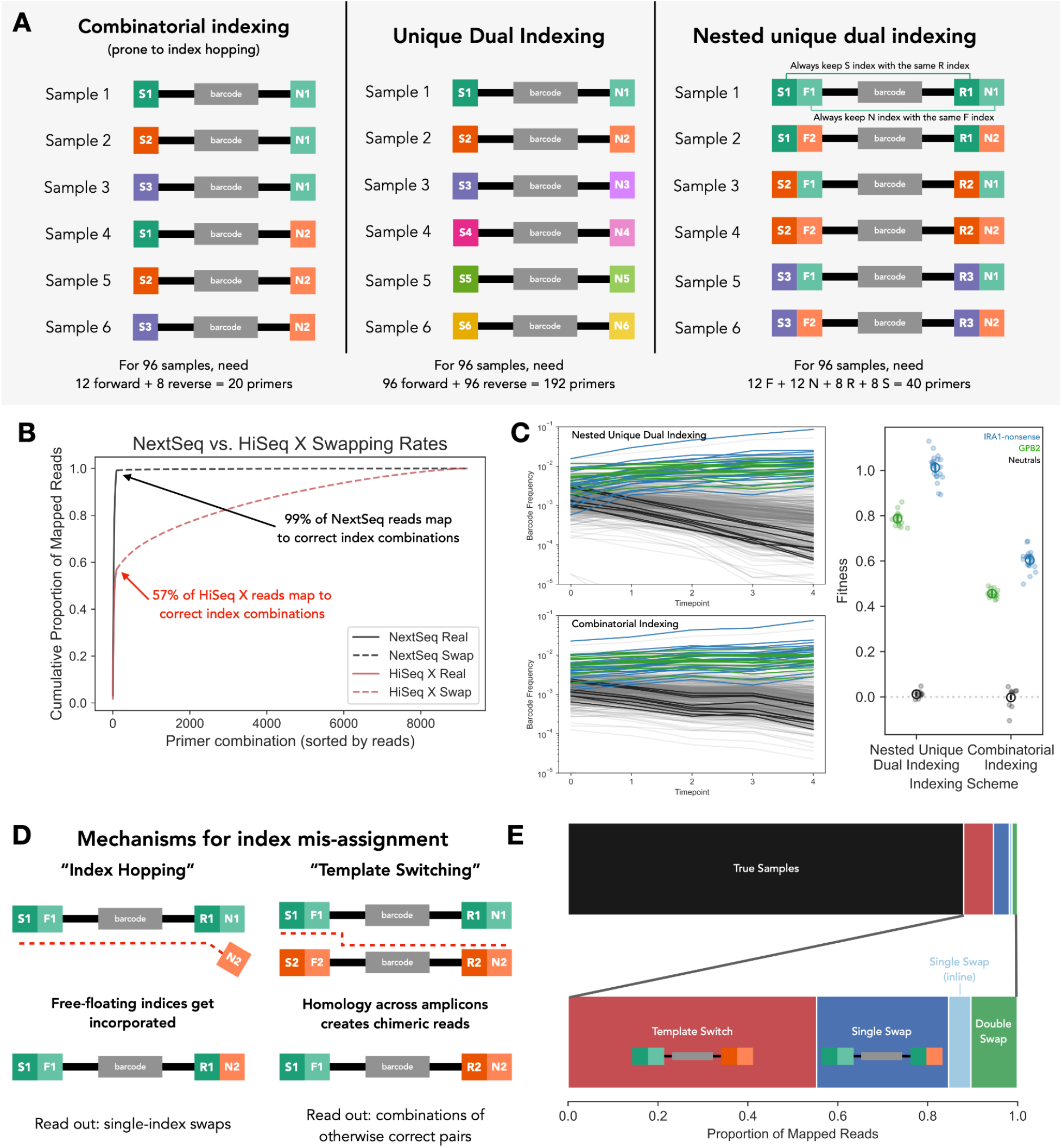
Template switching is a major source of incorrect sample assignment for barcode sequencing. **(A)** This panel shows three schemes for multiplexing samples onto the same sequencing lane. The first, “combinatorial indexing” assigns samples to many combinations of primers, allowing for many samples to be labeled with a relatively small number of indices. This scheme allows for index hopping to incorrectly assign samples. There are several alternatives that reduce the effect of index hopping and allow researchers to remove incorrect swaps from data. The most common scheme is “unique dual indexing”, where each sample uses a unique combination of forward and reverse indices. In this case, 192 primers are needed to label 96 samples. Another possible scheme, used in this paper, is nested unique dual indexing, where inline and Illumina indices are combined such that each inline index corresponds with a unique Illumina index on the other side of the amplicon molecule. This allows you to achieve unique dual indexing of 96 samples with 40 total primers. **(B)** Rates of incorrect sample index assignment are much higher on a HiSeq X (patterned flow cell) sequencing machine than NextSeq (non-patterned flow cell) for this set of 95 samples. **(C)** Undetected index mis-assignment can alter frequency trajectories and result in incorrect fitness inference. When index mis-assignment is detected using unique dual indexing (top left panel), frequency trajectories show a more dramatic change in frequency, especially for neutral lineages (black lines). In contrast, if combinatorial indexing were to be used, index mis-assignment would fail to be detected, resulting in diminished changes in barcode frequency over time (bottom left panel). The right panel shows the fitness values inferred under each scenario. The colored dots denote the inferred fitness for each barcoded mutant. The open circle shows the mean across all mutants for a particular genotype, with the error bar denoting standard error. **(D)** There are two primary mechanisms for index mis-assignment. One mechanism (depicted on left) is “index hopping” due to mis-incorporation of free-floating index primers present in the library. This results in reads that differ by a single index from correct index combinations. Another mechanism (depicted on right) is “template switching” due to switching of the template molecule during the amplification step of sequencing. This results in chimeric reads, where each end of the molecule matches a correct index combination but do not belong together. **(E)** For a library of 8 samples utilizing a nested unique dual indexing scheme consisting of 32 individual indices (such that each index was only used once), we can analyze which mechanisms caused index mis-assignment by looking at whether incorrectly indexed reads have one or more misplaced indices. The top stacked bar plot shows the proportion of reads where there was no index mis-assignment (black) and the proportion where indices were mis-assigned (colored). The bottom stacked bar plot shows the proportion of reads with mis-assigned indices that belong to each category of index hopping. We find that template switching (red) is the most commonly observed mechanism of index hopping, followed by single index swaps of Illumina indices (dark blue) and double swaps (green). Single index swaps of inline indices (light blue) occur at extremely low rates.

In order to quantify the extent to which this mis-assignment (often denoted as “index hopping”) affects amplicon libraries, we devised a nested dual-indexing approach that combines the use of inline indices and Illumina index primers. Specifically, we carry out two-steps of PCR, the first of which attaches inline indices (denoted by F and R in **Figure 3A**) to each side of the amplicon, followed by a second step which attaches Illumina index primers (denoted by N and S in **Figure 3A**) which also contain the P5 and P7 sequences which bind to the flow cell. In this scheme, we ensure that each inline index is only paired with a specific Illumina index on the other side of the molecule (i.e., R1 is always paired with S1, F1 with N1, R2 with S2, etc.) (**Figure 3A**). This allows us to detect index hopping events when one or more of the inline indices do not match their paired Illumina index.

First, we ran a single pooled library of 95 samples (consisting of all allowed combinations of primers except one) on one lane of NextSeq and one of HiSeq X. After demultiplexing the reads and mapping the reads to the template sequence, we find that only 1% of mapped NextSeq reads were assigned to incorrect combinations of indices. In contrast, 43% of mapped HiSeq X reads were assigned to incorrect combinations of indices (**Fig 3B**). This is much larger than the previously reported value of 5-10% of reads misassigned for diverse RNASeq or WGS libraries (Illumina; Sinha et al.). These very high rates of index mis-assignment can result in substantial effects on fitness measurement experiments. For example, if reads from a late timepoint are mis-assigned as coming from an early time point, lineages with high fitness (and thus high frequency in later time points), can have underestimated fitness effects, because their early time point frequency will be overestimated (**Fig 3C**). Additionally, high rates index mis-assignment can result in frequency trajectories that “zig-zag”instead of reflecting constant frequency change overtime, resulting in very noisy fitness estimates (KerryGeilerSamerotte 2018d; Kinsler 2018b).

Why is it that our barcode libraries have much higher rates of index hopping than previous reports focusing on RNAseq? There are two main mechanisms by which index mis-assignment could occur. First, free-floating indices present in the library could become incorporated into molecules initially tagged with another index, resulting in index-hopping (**Fig 3D**; left). Our experimental design can distinguish this type of index-hopping because this process should primarily occur through swapping of the Illumina index primers (which contain the flowcell-binding sequences; S and N in **Fig 3A** rightmost panel), rather than inline indices (F and R in **Fig 3A** rightmost panel). Therefore, we can identify these events by observing reads where all of the indices match an included sample except a single S or N Illumina index - we will refer to these events as “single swaps”. A second mechanism is that index mis-assignment occurs via “template switching events, where the polymerase jumps between two homologous sequences to create chimeric sequences where each end is from distinct original molecules (**Fig 3D**; right). Given that the diverse barcodes sequenced in amplicon libraries often include a homologous region that is identical between all molecules (Levy et al. 2015; Wong et al. 2016; Adamson et al. 2016; Najm et al. 2017; Gordon et al. 2020), this mechanism of swapping could represent a more severe problem for barcoders. In the case of our experimental design to detect index mis-assignment, if the swap occurs in the homologous region between the indices, we would expect to find mismatched index pairs. In other words, the index pair on one end of the molecule (i.e., S and F) would match one sample while index pairs on the other end of the molecule (R and N) would match a different sample (**Fig 3D**).

To distinguish between these two possible mechanisms, we deeply sequenced a library of 8 samples that were uniquely indexed using our nested unique dual index scheme (**Fig 3A**), such that each individual index was used with only a single sample (Kinsler 2018c). Sequencing this pooled library on a HiSeq X machine, we found that 88% of our mapped reads had correct combinations of indices (**Fig 3E**). Of the 12% remaining mis-assigned reads, 55% (6.6% of the total mapped reads) were likely products of template switching events, while 29% of the mis-assigned reads (3.5% of total mapped reads) were the product of single swaps of an Illumina index. In addition to these two prominent swapping events, 1.2% of total reads were double swaps (likely reflecting either multiple independent swapping events resulting in incorrect index combinations or residual contamination of some kind) and 0.5% of the total mapped reads were single swaps of an inline index.

These data suggest that template switching (**Fig 3D**; right) is a dominant driver of index mis-assignment when sequencing barcode libraries and could explain why mis-assignment rates are higher for amplicon sequencing than sequencing more diverse libraries, e.g., RNASeq or whole genomes. This makes sense given template switching relies on shared regions of homology between swapped reads, which are prevalent in amplicon sequencing (**Box 3**). This finding reinforces our recommended best practice of using unique dual indexing (or nested unique dual indexing) for barcode libraries, even in cases where you are confident there are few unincorporated primers remaining in the library, as these cause index-hopping (**Fig 3D**; left) but not template switching (**Fig 3D**; right).

#### Box 3 - The consequences of template switching on barcoded competition experiments

Given that template switching seems to occur more in barcoded libraries than in RNASeq or WGS data, we suspect that it is facilitated by homology amongst reads. Because of how barcode regions are synthesized and cloned, there are often fairly long stretches of homology flanking the barcode region. Thus, one potential area of future improvement is to consider barcoding designs that reduce the amount of homology between different barcodes (Hegde et al. 2018). Doing so is important not only to avoid wasting sequencing reads on swapped samples, but also to minimize swapping in cases where unique dual indexing cannot detect it. While unique dual indexing allows removal of template switching events between timepoints or samples, it does not detect swapping events between within the same sample. These “hidden” swapping events result in chimeric reads and can cause issues for certain applications. For example, in some methods including massively parallel reporter assays and barcoded or combinatorial CRISPR screens, it is important to associate information from one side of the read with the other (**Fig B3**) (Wong et al. 2016; Adamson et al. 2016; Najm et al.2017; Gordon et al. 2020). Researchers carrying out experiments that rely on these internal associations should utilize sequencing platforms with reduced template switching rates (e.g. Illumina non-patterned flow cell machines like NextSeq or MiSeq) and/or experiment with reducing homology amongst reads.

**Figure B3.**
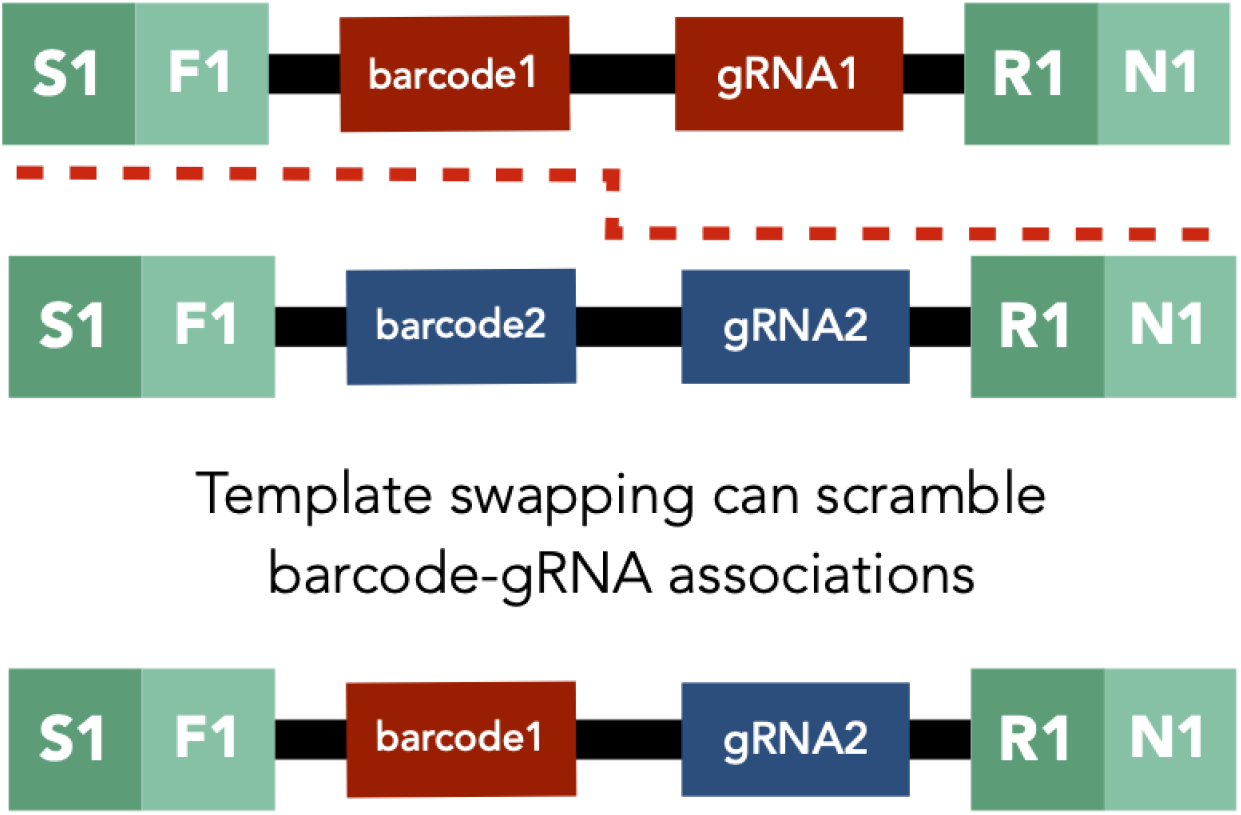
Template switching can scramble internal associations on different ends of the read. This figure shows a schematic of a template switching event occurring between two reads belonging to the same sample for an application where barcodes are associated with guide RNAs (i.e., barcoded CRISPR screen). In this case, template switching (denoted by the red dashed line) that occurs in the homologous region in the middle of the read would cause the gRNA-barcode associations to be scrambled. This would not be identified as a swapping event using unique dual indices because both reads belong to the same sample and have the same indices.

An important note about detecting index mis-assignment is that experiments with more samples will have more power to observe mis-assignment between samples. The reason being that we are blind to cases where swapping occurs within a sample, i.e., between molecules with the same set of indices. This likely underlies our observation of only 12% swapping events in an experiment with 8 samples (**Fig 3E**), but 43% in an experiment with 95 samples (**Fig 3C**). The previous study finds mis-assignment of 5 – 10% of reads in RNAseq data utilized 190 samples (Sinha et al.). Thus, despite being highly powered to detect swapping events between samples, it found far fewer than we find in barcode sequencing data, likely because template switching (**Fig 3D**; right) is more prevalent in amplicon sequencing. Though harder to detect, swapping within samples can represent a major problem (**Box 3**).

## Discussion

Here, we conducted many barcoded fitness measurements under conditions that were designed to reduce measurement error by as much as possible. Despite these efforts, we observed substantial variation in our fitness measurements, especially between replicate experiments and batch experiments done on different days (**Fig 1**). We investigated possible sources of noise, and suspect that variation across batches and replicate experiments is likely partially due to environmental differences between experiments and partially attributed to our fitness inference procedure. We found that technical noise due to DNA extraction and PCR amplification had minimal contributions to measurement noise when they were performed via the methods we used (**Fig 2**). And we found that another technical source of noise, template switching between sequencing reads from different samples, can provide a major source of noise (**Fig 3**). We conclude by providing a detailed list of best practices to contend with all of these sources of variation (**Box 4**).

### Box 4 - Best practices for barcoded fitness competitions; lessons from #1BigBatch

I. **Reducing technical variation across fitness estimates:**

a. **Ensure high sequencing coverage to avoid sampling noise:** Our measurements suggest that sequencing coverage is a driver of reproducibility. We found the worst reproducibility (R^2 < 0.8 in **Fig 2C**) when sequencing coverage dipped below an average of 20 reads per barcode (red points in **Fig 2C**), and that we had consistently better reproducibility when we sequenced on average 600 reads or more per barcode (blue points in **Fig 2C**).
b. **Ensure high levels of reference strain:** A mutant’s change in frequency depends on its fitness and the mean fitness of the population, which in some cases is inferred relative to a reference strain (**Box 2**). It is best practice to include the reference strain at sufficiently high frequency to be measured well, as well as including many uniquely barcoded reference strains in order to benchmark fitness with high precision.
c. **Choose amplification and extraction methods to minimize sample prep noise:** We recommend using established methods of DNA extraction and PCR amplification that have been shown to mitigate noise, e.g., two-step PCR with unique molecular identifiers.
d. **Use unique dual indexing to avoid index mis-assignment:** Template switching during Illumina sequencing can result in reads being assigned to incorrect samples and errant fitness estimates (**Fig 3**). To avoid these biases, we recommend using unique dual indexing schemes, such that mis-assigned reads can be identified and discarded.
II. **Contending with biological variation across fitness estimates:**

a. **Design highly-controlled experiments:** Batch effects, stemming from various uncontrolled factors, can result in substantial variation in fitness across days, labs, and personnel. It is best practice to minimize batch effects, for example, by using the same base medium for control and treatment conditions, and to assess the strength of batch effects by including a control experiment in every batch.
b. **Use many barcodes per mutation:** This allows you to estimate the uncertainty in a mutation’s measurement by taking many independent measurements in the same vessel, rather than measuring its fitness across many different experiments which may introduce batch effects.
c. **Embrace and report biological variation:** While batch-to-batch variation is often seen as a nuisance, it is reflective of rampant context-dependency in biological systems. We argue for embracing this context-dependency by openly reporting the variation in outcome alongside known subtle differences between experiments (e.g. media components, location in incubator, flask brand, frequency distribution of strains). This would allow researchers to leverage this variability to better understand biological systems.

One of our most upsetting observations was that fitness varies so much across replicates and batches. Taken at face value, the observation that the fitness of an organism can change so much between experiments done by the same group under the same conditions using the same methods raises questions about our ability to obtain and interpret fitness measurements. This concern is relevant in many fields, including experimental evolution (Levy et al. 2015; Li et al.2019; Boyer et al. 2021; Aggeli et al. 2021; Bakerlee et al. 2021), deep mutational scanning (Fowler and Fields 2014), and high-throughput genetic engineering systems (Sharon et al.2018; Bakerlee et al. 2022).

One way to contend with this concern is by reducing comparisons across replicates and especially batches as much as possible. For example, the fitness of all mutations could be measured at the same time in the same flask to reduce batch effects. In cases where this is not possible, for example, when comparing across two or more different environments, all experiments could be performed in the same batch, with as many factors held constant as possible (incubator, flask shape, base media, etc.). Indeed, this is how we performed the #1BigBatch experiment that inspired this manuscript. But performing all experiments in one big batch is difficult, and further, does not address the issue of reproducibility across labs. Should scientists studying fitness consider investing in ultra precise scales for use during media prep, or incubators with tight temperature control to ensure that others can more accurately recreate their work? We hope that the results reported here, particularly the large amount of variation in fitness observed across batches, inspires deeper discussion of this important issue.

On the other hand, perhaps batch effects need not be reduced. If the variation in fitness across replicates and batches does not reflect imprecise measurements, and instead results from small environmental differences between experiments, perhaps this is a time to make lemonade from lemons. Perhaps we should consider fitness as a parameter that inherently varies and can never be fully replicated even in the same experiment. If we embrace this context-dependency and report this variation instead of merely averaging away this signal, we might begin to glean new insights about these sensitive systems.

The observation that fitness is sensitive to subtle environmental changes across batches contributes to a growing mountain of evidence that context dependency is pervasive in biological systems (Eguchi et al. 2019; Liu et al. 2020; Kinsler et al. 2020; Bakerlee et al. 2021). This rampant context-dependency raises many philosophical issues about how to define an ‘environment’. Should we think of each batch as a new environment, or each replicate? When we perform an evolution experiment, do we expose our evolving population to a new environment every time we transfer them into new media? Is the environment also defined by which other genotypes are present and at what frequencies? Stepping outside of laboratory experiments, how should we think of the fitness of an organism if it is extremely sensitive to the small changes in the environment that happen over the course of the day? In particular, should we re-evaluate the use of models in which an organism has a single selection coefficient and perhaps define a range of coefficients that pertain to the relevant range of environments it experiences?

In sum, improvements in our ability to measure fitness have allowed us to detect rampant context-dependency hiding in a high-replicate dataset; ironically high-replicate datasets are often generated to improve measurement precision rather than reveal the inherent imprecision. The revelation that fitness is inherently varied across replicate experiments inspires new goals for fitness measurements going forward. In addition to measuring fitness precisely by eliminating technical sources of noise, the community must think carefully about how to measure an extremely context-dependent parameter in a reproducible way, how to report precision on these measurements, and how to use these measurements to understand and model processes such as natural selection, which can detect fitness differences orders of magnitude smaller than those we observe in the lab and thus may be even more sensitive to small environmental fluctuations.

## Methods

### Fitness measurements

The 28 replicate fitness measurements presented in **figure 1** were described in a previous study in great detail (Kinsler et al. 2020). Briefly, growth competitions were set up between a pool of barcoded mutants and a reference strain. The change in the frequency of each barcode over time reflects the fitness of the adaptive mutant possessing that barcode relative to the reference strain. After a growth competition is complete, DNA was extracted from frozen samples and resuspended in Elution Buffer to a final concentration of 50 ng/μL for later use in PCR reactions. A two-step PCR was used to amplify the barcodes from the DNA. The first PCR cycle used primers with inline indices to label samples. Attaching unique indices to samples pertaining to different conditions or timepoints allows us to multiplex these samples on the same sequencing lane. Each primer also contained a Unique Molecular Identifier (UMI) which is used to determine if identical barcode sequences each represent yeast cells that were present at the time the sample was frozen, or a PCR amplification of the barcode from a single cell. The second step of PCR used standard Nextera XT Index v2 primers (Illumina #FC-131-2004) to further label samples representing different conditions and timepoints with unique identifiers that allow for multiplexing on the same sequencing lane. Samples were uniquely dual-indexed following the nested scheme in **figure 3**. Pooled samples were then sent to either Novogene (https://en.novogene.com/) or Admera Health (https://www.admerahealth.com/) for quality control (qPCR and either Bioanalyzer or TapeStation) and sequencing. Data were processed by first using the index tags to de-multiplex reads representing different conditions and timepoints. Then, reads were mapped to a known list of barcodes, PCR duplicates were removed using the UMIs, and the frequency of each barcode was measured at each time point. Fitness was inferred from changes in barcode frequency over time using a modified version of fitness assay python (https://github.com/barcoding-bfa/fitness-assay-python) available at https://github.com/grantkinsler/BarcodersGuide, along with the rest of the code used in this study.

### Technical replicates

To create technical replicates of the DNA extraction procedure, frozen cell samples from a given time point of a pooled fitness competition were divided in half prior to DNA extraction. The halved samples were then treated separately for all downstream steps. All sample preparation was conducted as described above and as detailed in Kinsler et al 2020, except in a few cases. In some of these exceptional cases, marked by a triangle in **figure 2C**, we omitted glass beads from one sample in the pair of technical replicates. In other exceptional cases, marked by a diamond in **figure 2C**, we added phenol to one sample in the pair of technical replicates. These modifications did not seem to affect reproducibility between the pair of samples (**Fig 2C**). Similarly, for technical replicates of the PCR procedure, we divided samples in half after DNA was extracted and diluted to 50 ng/μL. These samples were processed independently for all downstream steps, following identical procedures as described in Kinsler et al 2020 with a few exceptions. In some samples, marked by a circle in **figure 2C**, we reduced the cycle time for one sample in the pair of technical replicates from 27 to 23 cycles. This did not affect reproducibility between a pair of samples (**Fig 2C**). All technical replicates were performed on samples from the batch of pooled fitness competitions initiated on 12/10/17; this was the batch of experiments that we live tweeted about at #1BigBatch.

### Quantifying the effects of index hopping

Below, we describe the methods we used to index samples, identify index mis-assignments, and understand the mechanisms underlying index mis-assignment. Note that other than the section *Template switching on patterned flow cells appears to be a major source of noise,* results presented throughout the rest of the study were sequenced to minimize the amount of index mis-assignment as possible. In particular, we included only samples that were sequenced on non-patterned flow cell technology (HiSeq 2000, HiSeq 2500, or NextSeq) or were sequenced on patterned flow cell technology (patterned flow cell HiSeq X) with nested unique-dual indexing.

We performed nested unique-dual indexing following the method we developed in previous work (Kinsler et al. 2020). Briefly, this approach uses a combination of inline indices attached during the first step of PCR (12 forward and 8 reverse), as well as Illumina indices (12 Nextera i7 and 8 Nextera i5) attached during the second step of PCR. The latter indices are not part of the sequencing read (they are read in a separate Index Read). To uniquely label each sample,we use combinations of the Illumina and inline primers, such that each of the 12 forward inline (F) indices is used solely with one Nextera i7 (N) index and each of the 8 reverse inline (R) indices is used solely with one Nextera i5 (S) index. We can then combinatorially utilize these 12 F/N combinations with the 8 R/S combinations to uniquely label up to 96 samples (**Fig 3**).

To estimate the magnitude of index mis-assignment for a large set of samples across different sequencing platforms, we first constructed a library by pooling 95 samples that were uniquely dual indexed in this manner. We then sequenced this same library on two lanes, one a NextSeq lane and the other a HiSeq X lane. During the processing of the sequencing data, we mapped reads to all possible combinations of indices and counted the number of mapped reads to each index combination. Any reads which mapped to incorrect index combinations were then classified as mis-assigned.

To demonstrate an example of the effect that unidentified index hopping can have on frequency trajectories and the calculation fitness, we used samples from replicate D of the experiment conducted on 12/10/17. The results depicting nested dual indexing use the reads to these samples from the HiSeq X run. To show what would happen if only combinatorial indexing were used, we ignored the inline indices, and added up all the mapped reads to each barcode that mapped to the proper Illumina indices (N and S) alone. We then calculated fitness using these new reads.

To quantify whether index mis-assignment occurs most frequently via template switching or single swaps, we separately pooled a set of 8 samples, each containing their own unique index at all positions (except on particular S index - see note below), and sequenced this on a lane of HiSeq X. We then mapped these reads to every possible index combination. Reads were classified as a template switching event if the indices on both sides of the molecule matched proper samples but should not exist together (**Fig 3D**). Reads were classified as a single swap if all indices matched a true sample except one. If the incorrect index was an Illumina index (N or S), it was classified as a single Ilumina swap, otherwise, if the incorrect index was in the read, it was classified as an inline single swap. Finally, if two indices were incorrect but the index combination was not due to a template switching event in the middle of the molecule, it was classified as a double swap.

Note that during the library creation for these samples, S513 was incorrectly used instead of S517 for four samples and S517 was used instead of S513 for four samples. For analysis about the general rate of index mis-assignment for amplicons, this may slightly decrease our estimated rate of index hopping, as single swaps between S513 and S517 would not be detected for the samples where the index was incorrectly used (likewise, single inline swaps between the corresponding inline R301 and R304 indices that are supposed to be uniquely associated with these samples. Note that this only applies to a small number of the total combinations that can arise via index mis-assignment, so the impact of this error should be minor. Only one of these samples was included in the library used to identify the mechanisms underlying index mis-assignment for amplicons. This resulted in the S517 index represented twice in this sample rather than only once. This could result in a slight under-estimation of the number of single swap events with Illumina primers (as again a single swap between the S517 index of these two samples would not be distinguished). Additionally, this error results in two primer combinations which could be either the result of single inline swaps or template switching. As template switching events are over 10x more likely than single inline swaps according to our data, we assigned these ambiguous combinations as template switching events. Because these events are much more common and this only applies to 2 of the 56 possible combinations that result from template-switching, this should result in only a minor mis-estimation of the template-switching and inline single swap rates.

## Acknowledgements

This work was supported by a National Institutes of Health grant R35GM133674 (to KGS), an Alfred P Sloan Research Fellowship in Computational and Molecular Evolutionary Biology grant FG-2021-15705 (to KGS), a National Science Foundation Biological Integration Institution grant 2119963 (to KGS), a National Institutes of Health grant R35GM118165 (to DP), a Chan Zuckerberg Investigator Award (to DP), and a Stanford Center for Computational, Evolutionary, and Human Genomics predoctoral fellowship (to GK).

